# Ecological effects of stress drive bacterial evolvability under sub-inhibitory antibiotic treatments

**DOI:** 10.1101/2020.06.30.181099

**Authors:** Marie Vasse, Sebastian Bonhoeffer, Antoine Frenoy

## Abstract

Stress is thought to increase mutation rate and thus to accelerate evolution. In the context of antibiotic resistance, sub-inhibitory treatments could then lead to enhanced evolvability, thereby fueling the adaptation of pathogens. Conducting a meta-analysis of published experimental data as well as our own experiments, we found that the increase in mutation rates triggered by antibiotic treatments is often canceled out by reduced population size, resulting in no overall increase in genetic diversity. A careful analysis of the effect of ecological factors on genetic diversity revealed that the potential for regrowth during recovery phase after treatment plays a crucial role in evolvability and is the main predictor of increased genetic diversity in experimental data.

## 2 Main text

A key aspect of the fight against antibiotic resistance concerns the origin of the genetic innovations generating resistance phenotypes. It has been suggested that many abiotic stresses, including sub-inhibitory antibiotic treatments, can increase mutation rate [1– 5]. This implies that low doses of antibiotics would not only select for pre-existing resistance alleles [6], but could also accelerate the generation of random genetic diversity, including rare antibiotic resistance alleles [7]. The actual –realized– genetic diversity, however, is not only determined by mutation rate. Population dynamics, and population size in particular, further plays a crucial role by driving the number of individuals on which selection can act [8, 9]. In the context of antibiotic resistance, this is for example well reflected by the finding that the evolution of antibiotic tolerance often precedes resistance [10]. Therefore, understanding the role of antibiotics on the ability of a population to generate genetic diversity, *i*.*e*. on evolvability, critically relies on combining the study of the physiological effects of antibiotics at the molecular level (e.g. mutagenesis) with the study of their impacts on bacterial population dynamics [11].

Properly isolating and quantifying mutation rates, doubling times and death rates is not always possible. The estimation of mutation rate in particular brings intrinsic difficulties, because it relies on sophisticated mathematical models making strong biological assumptions (including no cell death) that are not always pertinent, especially under stress [11]. Instead, we suggest using a simple and easy-to-collect metric to quantify the effect of treatment on the generation of genetic diversity, and thus on evolvability (defined as the capacity of a population to generate adaptive genetic diversity, and therefore to evolve by natural selection). This metric directly relies on the observed number of mutants towards a neutral arbitrary phenotype. It is calculated as the ratio of the observed number of mutants in treated populations to the observed number of mutants in untreated controls (see Methods for details). While the observed number of mutants is not exactly the mutation supply, because the same mutation event can give several clones in the final population, both variables are closely related and the former can be directly measured in practice.

To assess the role of antibiotics on bacterial evolvability, we collected raw data from primary literature on the effect of antibiotic treatments on mutation rates, and conducted a meta-analysis with emphasis on the effect of treatments on both mutation rate and population size. Additionally, we performed our own set of experiments for the most commonly studied antibiotics, to mitigate the difficulty of comparing data from different labs in meta-analyses. We show that the decrease in population size due to the treatment often cancels the potential increase in mutation rate, resulting in no overall increase of evolvability. We further combine the experimental data with simulations to explore the ecological conditions in which antibiotic treatments may still increase the generation of genetic diversity.

Our work stems from the simple intuition that an antibiotic treatment which increases mutation rate by 10 fold but decreases population size by 100 fold is not likely to increase evolvability (as for example noted by Couce et al. [12]). We confirmed this intuition with simple stochastic simulations of the arisal of neutral mutations in a population subject to demographic forces (constant birth and death rates) with a constant mutation rate. We found that, in this simple scenario, the treatment decreases the generated genetic diversity (*i*.*e*. the total number of mutants and of unique mutational events, *p <* 10^−9^, Fig. 1). This is, for example, consistent with the vague definition of mutation supply as the overall product of mutation rate and population size (eg [13, 14], following a more precise definition of mutation supply per generation in the context of Moran’s model by Maynard Smith [15]).

**Figure 1:**
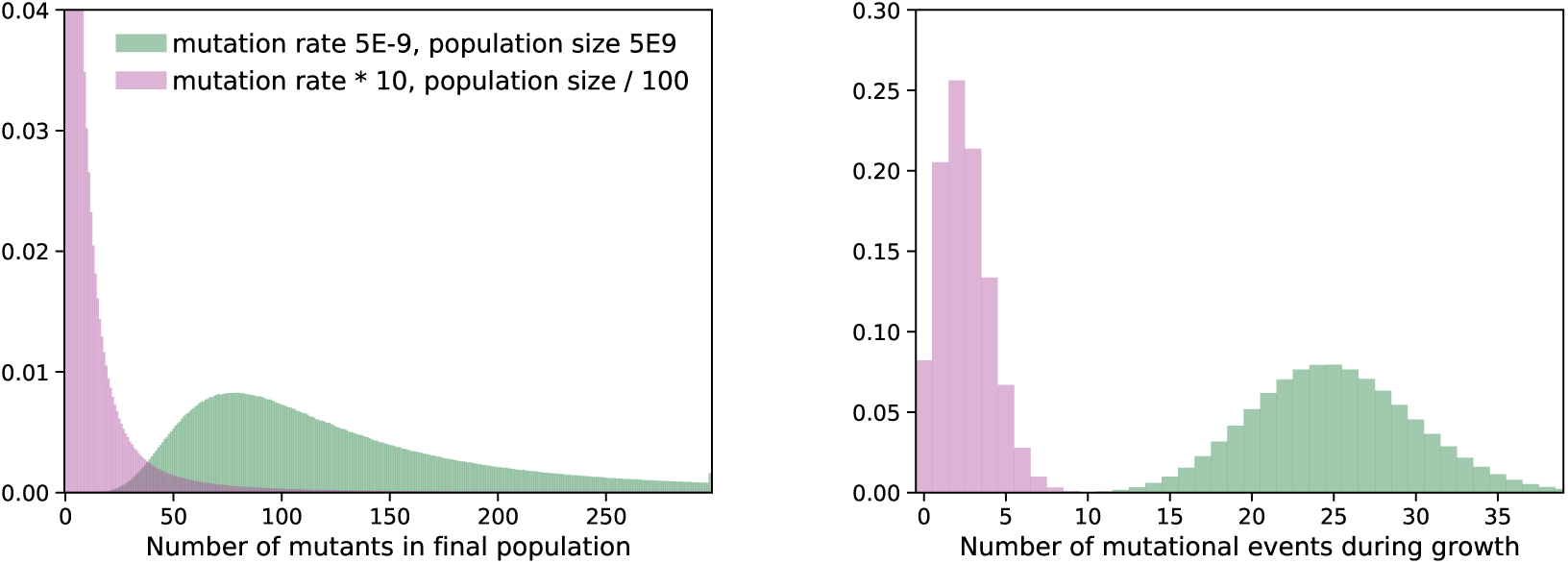
Change in genetic diversity due to a hypothetical treatment that increases mutation rate (x10) but decreases population size (/100). The left panel shows the distribution of the number of mutants in the final population for 10,000,000 simulations. The right panel shows the distribution of the number of mutational events in the same simulations. The number of mutational events gives a more precise estimation of genetic diversity, because a single mutational event can give several times the same mutant in the final population. The number of (non-unique) mutants, however, can be directly measured experimentally.

This observation questions whether the antibiotic treatments reported in the literature to increase mutation rate also have an effect on population size, and whether this effect may outbalance the reported increase in mutation rate, resulting in unchanged or even decreased genetic diversity. To address this question, we re-analyzed the raw data from 10 published papers on the effect of sub-inhibitory antibiotic treatments on mutation rate (Table 1) [11, 16–24], that we complemented with our experimental data. We first evaluated the effect of the antibiotic treatments on both computed mutation rates and population sizes. Estimating mutation rates with a widely used modern method (rSalvador [25]), we confirmed a systematic increase in computed mutation rates in the presence of sub-inhibitory doses of antibiotics in both data from the literature and our own data (average increase over all data points: 4.49 fold, significantly higher than 1, *p* = 7.53 × 10^−10^, Fig. 2A). However, at these concentrations reported to increase mutation rate, antibiotic treatments overall concomitantly decreased population size (*p* = 1.08 × 10^−5^, Fig. 2B).

**Table 1:**
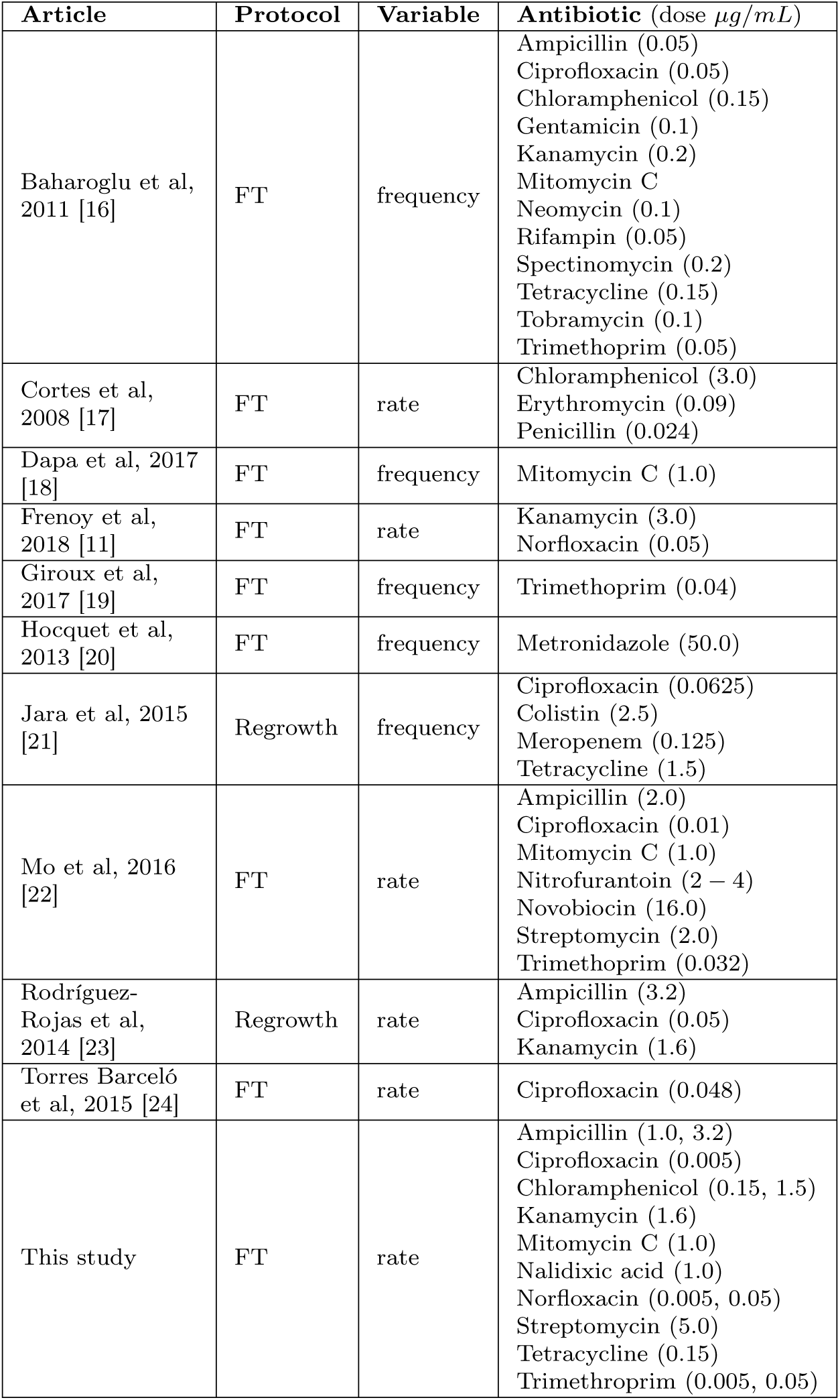
Experimental data analyzed in this work: origin, protocol, reported variable and antibiotic treatment. The protocol can be ‘FT’, indicating a standard fluctuation test, or ‘Regrowth’, indicating that cultures recovered in fresh antibiotic-free medium following antibiotic exposure. The reported variable indicates whether the original study reported mutation rates or mutation frequencies. In both cases, we use the raw data for mutant counts, population size and plating fraction to analyze all datasets with the same methods.

| Article | Protocol | Variable | Antibiotic (dose $\mu\text{g/mL}$) |
| --- | --- | --- | --- |
| Baharoglu et al, 2011 [16] | FT | frequency | Ampicillin (0.05)<br>Ciprofloxacin (0.05)<br>Chloramphenicol (0.15)<br>Gentamicin (0.1)<br>Kanamycin (0.2)<br>Mitomycin C<br>Neomycin (0.1)<br>Rifampin (0.05)<br>Spectinomycin (0.2)<br>Tetracycline (0.15)<br>Tobramycin (0.1)<br>Trimethoprim (0.05) |
| Cortes et al, 2008 [17] | FT | rate | Chloramphenicol (3.0)<br>Erythromycin (0.09)<br>Penicillin (0.024) |
| Dapa et al, 2017 [18] | FT | frequency | Mitomycin C (1.0) |
| Frenoy et al, 2018 [11] | FT | rate | Kanamycin (3.0)<br>Norfloxacin (0.05) |
| Giroux et al, 2017 [19] | FT | frequency | Trimethoprim (0.04) |
| Hocquet et al, 2013 [20] | FT | frequency | Metronidazole (50.0) |
| Jara et al, 2015 [21] | Regrowth | frequency | Ciprofloxacin (0.0625)<br>Colistin (2.5)<br>Meropenem (0.125)<br>Tetracycline (1.5) |
| Mo et al, 2016 [22] | FT | rate | Ampicillin (2.0)<br>Ciprofloxacin (0.01)<br>Mitomycin C (1.0)<br>Nitrofurantoin (2 – 4)<br>Novobiocin (16.0)<br>Streptomycin (2.0)<br>Trimethoprim (0.032) |
| Rodríguez-Rojas et al, 2014 [23] | Regrowth | rate | Ampicillin (3.2)<br>Ciprofloxacin (0.05)<br>Kanamycin (1.6) |
| Torres Barceló et al, 2015 [24] | FT | rate | Ciprofloxacin (0.048) |
| This study | FT | rate | Ampicillin (1.0, 3.2)<br>Ciprofloxacin (0.005)<br>Chloramphenicol (0.15, 1.5)<br>Kanamycin (1.6)<br>Mitomycin C (1.0)<br>Nalidixic acid (1.0)<br>Norfloxacin (0.005, 0.05)<br>Streptomycin (5.0)<br>Tetracycline (0.15)<br>Trimethoprim (0.005, 0.05) |

**Figure 2:**
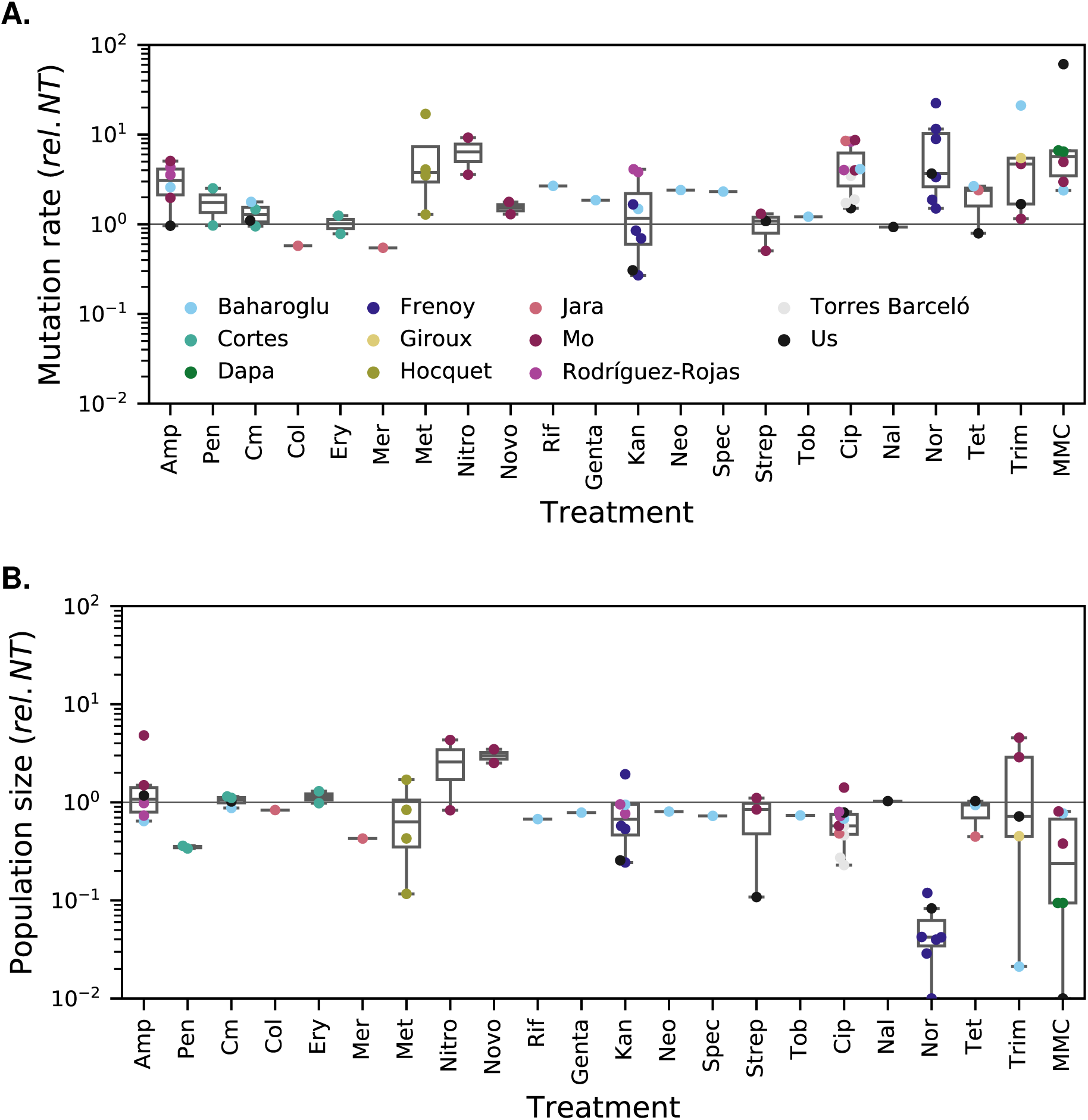
Effect of antibiotic treatments on estimated mutation rate and population size. Different antibiotics are shown in different columns on the x axis, ordered by class of action. Colors indicate different studies (10 datasets from the literature plus our data). A single study may comprise several data points associated with the same antibiotic because several doses were tested or because several biological replicates (each of them comprising several parallel replicate populations) were performed. **A. Effect of antibiotic treatments on estimated mutation rate**, under the assumption that there is no cell death. All mutation rates are relative to the untreated control (rel. NT, horizontal line y = 1). The mutation rates were estimated using rSalvador [25]. **B. Effect of antibiotic treatments on population size**. All population sizes are relative to the untreated control (horizontal line y = 1). Each data point represents the average final population size in several parallel replicate populations treated with a specific dose of a given antibiotic.

We then addressed the central question of the overall outcome of decreased population size and increased mutation rate on genetic diversity. Using our proposed metric of evolvability, we found that the effect of antibiotic treatments on the number of mutants for the neutral phenotype of interest is more equivocal, with great variation both between and among antibiotics (Fig. 3). Interestingly, the data points indicating higher evolvability, *i*.*e*. more mutants compared to the untreated baseline, almost all belonged to a small subset of studies. We therefore wondered whether systematic differences between the experimental conditions could account for this finding. We found that the studies in which genetic diversity increased were (1) those which used a variant of the fluctuation test protocol in which populations recover in fresh medium after antibiotic exposure, resulting in regrowth after treatment, and (2) the one study in which population size was found to increase for several treatments.

**Figure 3:**
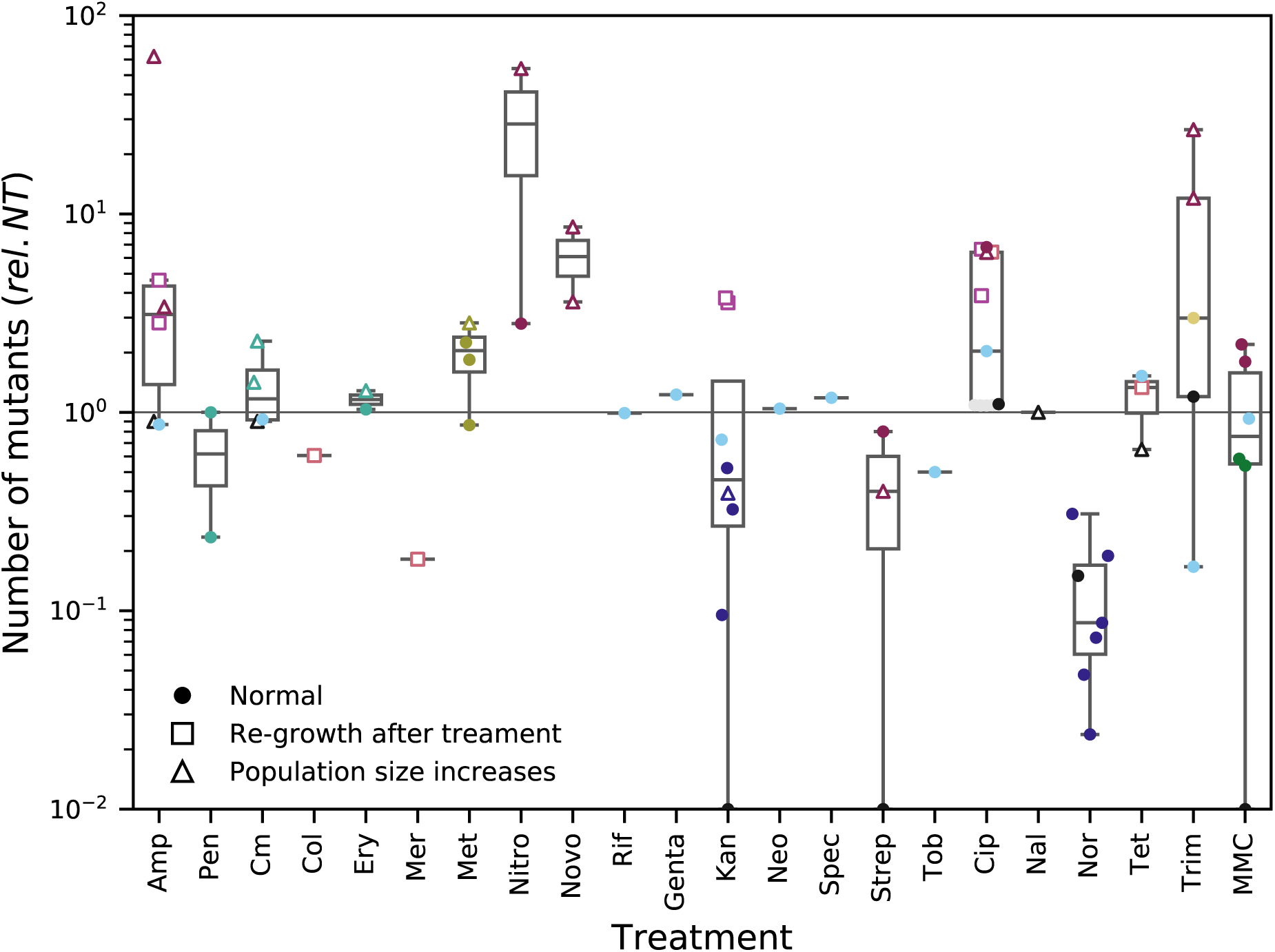
Effect of antibiotic treatments on genetic diversity. The raw number of mutants on selective medium is used as an approximation of genetic diversity. This number of mutants is relative to the untreated control (horizontal line y = 1). The colors correspond to different studies and are the same than on Fig. 2. Open squares indicate the use of a modified fluctuation assay protocol where treated cells are regrown in fresh antibiotic-free medium before being plated. Open triangles indicate the particular situation where antibiotic treatment increases population size. All other data points (standard fluctuation assay, no increase in population size) are labeled with filled circles.

Integrating these factors into our analysis of the effect of treatment on genetic variability, we found that data points associated with increased population size or with regrowth after treatment have a significantly increased evolvability relative to untreated (mean relative evolvability 7.95, significantly higher than 1, *p* = 0.0011). Conversely, data points from standard fluctuation assay protocol show no detectable increase in evolvability (mean relative evolvability 1.03, no evidence that higher than 1, *p* = 0.13). The difference between the evolvability of the data points from the two categories is highly significant (*p* = 8.21 × 10^−5^).

The effect of regrowth after exposure to antibiotic stress can be easily understood, in the light of the intuition that population size is a key driver of genetic diversity. Simulations of treatments with and without regrowth confirm the effect detected in the meta-analysis (Fig. S2, S3): genetic diversity is increased (both compared to untreated and treatment without regrowth) for a treatment followed by a regrowth phase. Regrowth should however not be merely regarded as a peculiarity of this protocol (that is designed to let the cells recover from filamentation [26]). Indeed, antibiotic treatments are frequently heterogeneous in time and space, and pathogens as well as commensals in a complex biotic environment such as the gut microbiome may thus be subject to cycles of treatment and regrowth. In such cases, the combination of increased mutation rates and higher population turnover and size could lead to greater genetic diversity, with potential consequences for the evolution of antibiotic resistance.

Our finding that increase in mutation rates is often compensated by a decrease in population size could be analyzed in the context of the drift-barrier hypothesis for the evolution of mutation rate [27]. This hypothesis suggests that DNA replication is always selected for highest accuracy (consistently with the reduction principle [28]), but that effective population size is a barrier to the efficiency of such selection. As a consequence, mutation rate is predicted to be inversely correlated with population size among microbial species [27]. While error-prone polymerases [29, 30] are often thought to be selected to generate more diversity under stress [31, 32], in the context of second-order selection for evolvability [33, 34], a similar alternative reasoning could hold here: because these polymerases repair DNA damaged by various exogenous stresses, they could systematically act under conditions of reduced effective population size and higher genetic drift (random death), and thus not be as efficiently selected for accuracy.

With mutation rate computation posing many theoretical challenges, we propose to estimate evolvability using a simpler and directly measured variable, *i*.*e*. the raw number of observed mutants for an arbitrary selectable phenotype in the treated population relative to the untreated controls. We conclude that at the chosen concentrations, antibiotic treatments significantly decreased genetic diversity when the bacteria are not regrown after treatments. While mutation rate is a crucial variable for the understanding of the molecular effects of various stresses, we show here that it is a poor predictor of evolvability when stress affects population demographics.

## 4 Acknowledgments

We express our deepest gratitude to all the scientists who shared their raw data with us, digging in their archives sometimes more than ten years after publication: Jesús Aranda Rodríguez, Zeynep Baharoglu, Tanja Dapa, Xavier Giroux, José Echenique, Arnaud Gutierrez, Didier Hocquet, Charlie Mo, Alexandro Rodríguez-Rojas, Clara Torres Barcelo, Christiane Wolz, Denise O’Sullivan, Timothy McHugh, Christiane Wolz, and their collaborators.

## Funding

This work was supported by the ERC (Grant number 268540, FP7-IDEAS-ERC program, attributed to S.B.), SNF (Grant number 155866, attributed to S.B.), and Marie Curie-ETH cofund Fellowship (Grant number 16-2 FEL-59, attributed to M.V.).

## Author contributions

M.V. and A.F. performed the experiments and analyzed the data. A.F., M.V. and S.B. conceived the research and wrote the paper.

## Competing interests

The authors declare no competing interests.

## 5 Supplementary materials

### Material and methods

#### Meta-analysis of previous studies

We focused on studies posterior to 2008 (no more than 10 years old at the beginning of this project) which quantified the effect of sub-inhibitory antibiotic treatments on bacterial mutagenesis. We were able to gather raw data for 10 studies (see Table 1). For each study, we extracted the raw data of the number of mutants (toward a chosen neutral phenotype) and total bacterial densities both after antibiotic exposure and in untreated controls, and the fraction of population plated on selective medium to score mutants. This list of variables reflects important features of mutation rate estimation using the standard fluctuation test devised by S. E. Lüria and M. Delbrück: the initial bacterial density of the founder population does not matter as long as it is small, and partial plating needs to be mathematically accounted for [35].

The following studies contain relevant data, but were excluded from the analysis despite matching our *a priori* inclusion criteria: Gutierrez et al. [5], Schroder et al. [36] (data available, but in a different representation or in a summary form); Kohanski et al. [4], Nair et al. [37], Peng et al. [38], Bunnell et al. [39], Cairns et al. [40], Thi et al. [41], Song et al. [42], Valencia et al. [43], Nagel et al. [44] (raw data unavailable or no answer from the author who kept the raw data).

#### Alternative metric of evolvability

In our attempt to integrate both mutation rate and population dynamics in evolvability, we propose to use the ratio of raw number of mutants towards the neutral phenotype of interest in treated populations to raw number of mutants in untreated controls. This metric has the additional advantage to be easy to collect in the lab. However, due to the large effect of the timing of appearance of mutations on the final number of mutants in the fluctuation test, this method requires a relatively high level of replication (compared to traditional binomial or poissonian dilution-sampling-plating assays). This is the case for all methods derived from the fluctuation test.

The reproduction of mutants that emerged early in the culture also means that each observed mutants in the final population does not necessarily correspond to a unique mutational event and thus to a unique genotype. However, as seen on Fig. 1, the number of unique mutational events is closely related to the number of observed mutants, and antibiotic treatment is unlikely to systematically bias the relationship between both variables.

This allows us to extrapolate the observed number of mutants towards an arbitrary phenotype (one that can easily be selected by plating such as the traditionally used rifampicin resistance conferred by mutations in rpoB) to a general measure of genetic diversity in the population. The underlying assumption for this generalisation is that mutation rate is increased in a similar fashion at all positions on the genome, although the mutation spectrum may be affected by stress [45].

#### Experimental detection of *de novo* mutations under sub-inhibitory antibiotic treatments

We performed a protocol closely inspired by the historical fluctuation test [46]. The aim was not only to add new data for the most studied antibiotics, but also to generate data for different antibiotics under strictly comparable conditions.

#### Culturing conditions

Using an overnight culture of *Escherichia coli* MG1655, we applied a strong bottleneck to initiate 12 parallel replicate populations. Specifically, we diluted the culture 10^5^ times and inoculated 10 *µ*L into 1 mL of fresh LB (lysogeny broth, Miller formulation) supplemented or not with antibiotics, in 13 mL tubes (Sarstedt, Germany). We used 20 antibiotic conditions with 12 replicates each for a total of 240 cultures (9 different antibiotics with several doses and untreated control, see Table 1). We incubated the cultures at 37°C shaken at 300 rpm for 24 hours.

#### Estimation of mutant frequencies

After 24 hours of growth in the presence of the different antibiotic treatments, we estimated the number of *de novo* mutants by plating 200 *µ*L of each culture onto LB agar supplemented with 100 *µ*g/mL rifampicin (*i*.*e*. rifampicin resistance is the neutral phenotype chosen to score *de novo* mutations). We further plated serial dilutions of six randomly chosen populations for each treatment onto LB agar to estimate total bacterial densities.

#### Mutation rate

We estimated mutation rate in the datasets from the literature and in our own experiments using Rsalvador v1.7 [25], with a correction for partial plating. This estimation does not account for potential bacterial death during the experiment.

#### Simulations

Simulation data presented on Fig. 1, S1, S2, S3 were obtained using adaptive tau-leap simulation of accumulation of a neutral mutant allele in a population subject to known constant demographic forces (growth and death with maximal carrying capacity) and mutational forces (mutation rate per individual per division toward a neutral mutant genotype).

For the presented simulations on Fig. 1, death rate is null, mimicking a purely bacteriostatic action of the antibiotic. Fig. S1 presents the results of similar simulations, but with an antibiotic that also has a bactericidal action (death rate 0.5 relative to birth rate).

For each figure and each condition, 10, 000, 000 replicate simulations were performed. The simulation program is included in the data deposit.

## Supplementary figures

**Figure S1:**
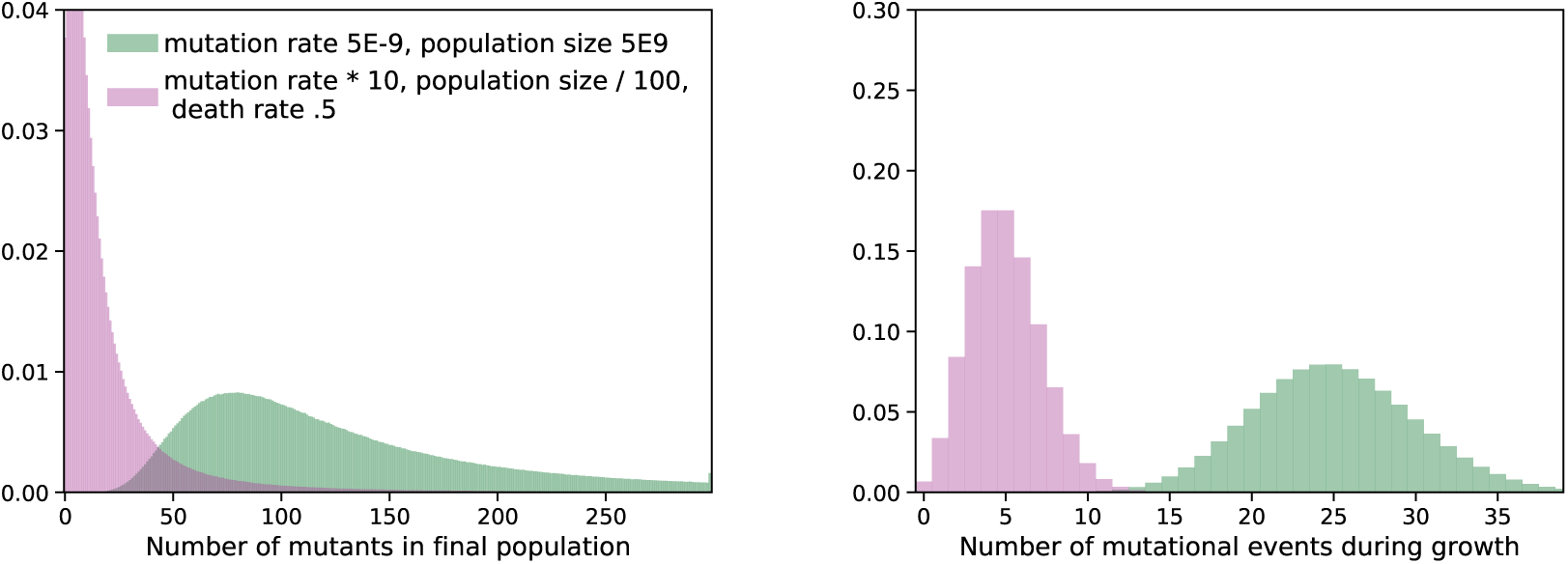
Change in genetic diversity due to a hypothetical treatment that increases mutation rate (x10) but decreases population size (/100), with death rate 0.5. This is similar to figure 1, but with a treatment that has a bactericidal activity and not only a bacteriostatic one.

**Figure S2:**
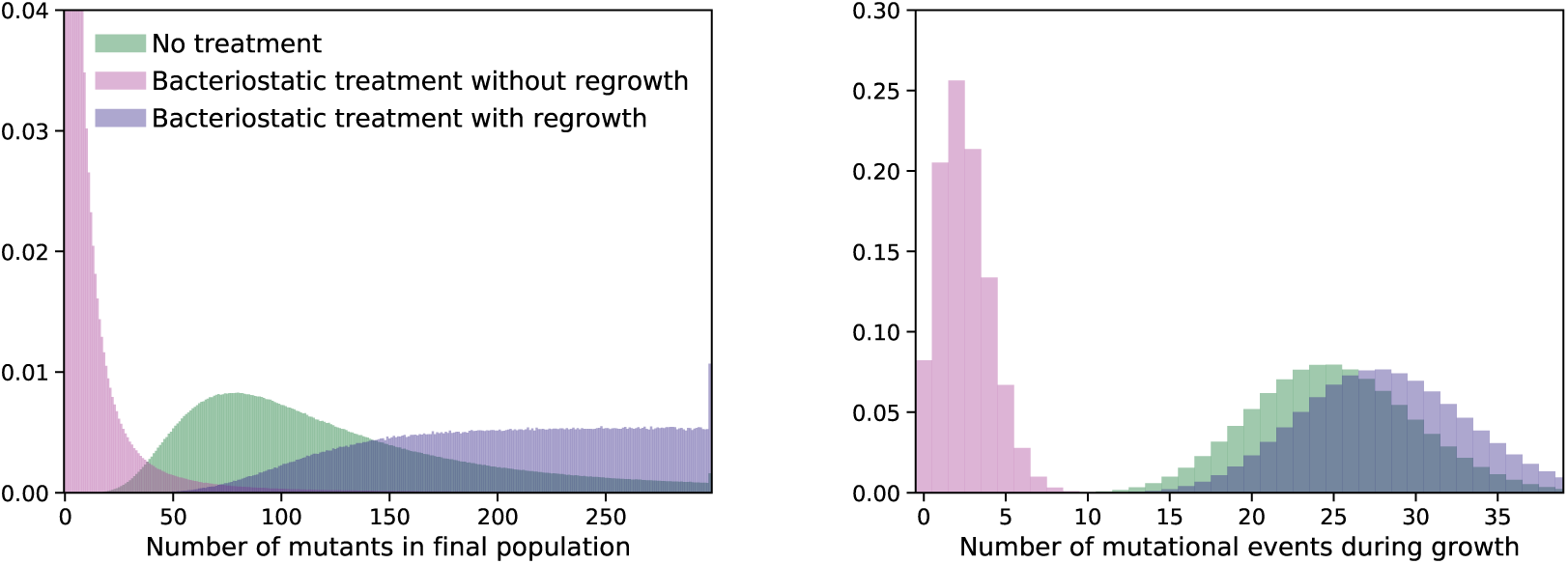
Change in genetic diversity with and without regrowth for a bacteriostatic treatment that increases mutation rate (x10) but decreases population size (/100). The data for untreated population and treatment without regrowth are the same than on Fig.1. The data for regrowth are produced by similar simulations, but with two phases: treatment (mutation rate x10) followed by recovery (mutation rate back to normal).

**Figure S3:**
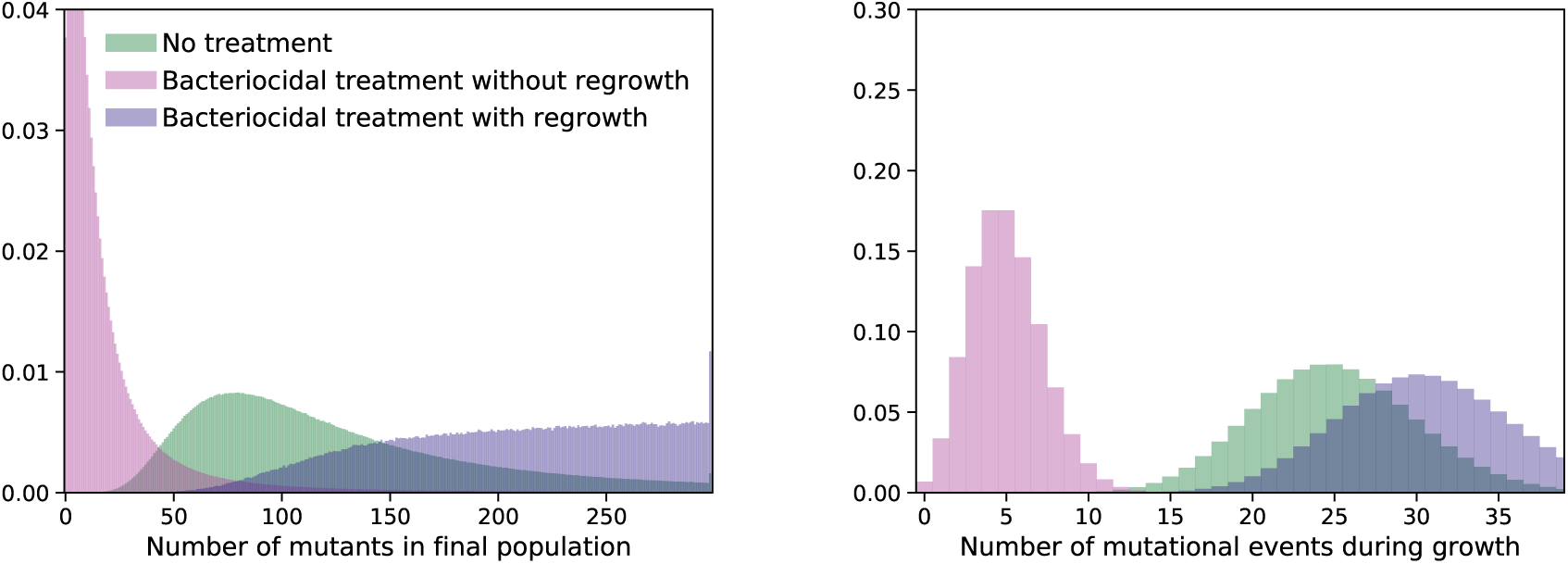
Change in genetic diversity with and without regrowth for a bactericidal treatment that increases mutation rate (x10) but decreases population size (/100) with death rate 0.5. This is similar to figure S2, but with a treatment that has a bactericidal activity and not only a bacteriostatic one. The relationship between the number of mutational events during growth and the number of mutants in the final population is more complex, because the lineage of a mutant may get extinct.

## Notes

### Competing Interest Statement

The authors have declared no competing interest.

